# The role of a G-quadruplex structure in mediating the interactions of miR-1249-3p with the 3’-UTR of the *HNRNPK* mRNA

**DOI:** 10.64898/2026.10.09.758038

**Authors:** Manju Kasaju, Rosalia M. Palumbo, Carlan Gray, Lydia Grimaldi, Jacob Shoulders, David B. Heisler, Mihaela-Rita Mihailescu

## Abstract

Heterogenous nuclear riboprotein K (hnRNP K) is an important regulator of multiple signaling pathways involved in DNA repair and function whose expression has been found to be mis-regulated in many cancers and neurological diseases. The expression regulation of hnRNP K is not fully understood however it has been shown that the microRNA miR-1249-3p targets its mRNA 3’-untranslated region (UTR), regulating its translation. In this study we identified a guanosine-rich sequence in the *HNRNPK* 3’-UTR within the miR-1249-3p binding site that has the potential to form a G-quadruplex (GQ) structure. By using various biophysical methods, we show that this sequence forms a parallel GQ structure which modulates the binding of miR-1249-3p to *HNRNPK* 3’-UTR *in vitro*. Additionally, using luciferase reporter assays, we determined that this GQ structure affects the hnRNP K protein expression, suggesting modulation of the binding interaction of miR-1249-3p to the 3’-UTR of *HNRNPK*. This *HNRNPK* mRNA GQ-miR binding axis is an unexplored mechanism of regulation, offering a novel therapeutic target for diseases caused by abnormal hnRNP K expression.

## Introduction

The heterogeneous nuclear ribonucleoprotein (hnRNP) family consists of multifunctional and multifaceted proteins that play important roles in nucleic acid biogenesis.^1^ Proteins in the hnRNP family, categorized as hnRNP A1 to hnRNP U, have at least one of the four unique RNA binding domains (RBDs): the RNA recognition motif (RRM), quasi-RRM, K-homology domain (KH), and glycine rich RNA binding domain.^2^ Among these proteins, hnRNP K is structurally and functionally distinct due to the presence of one additional unique K-protein interactive (KI) domain.^3^

hnRNP K has many different roles in the nucleus and cytoplasm, mainly acting as a protein hub to transport multiple proteins and RNAs that directly or indirectly influence signaling pathways.^4^ Altered expression of hnRNP K has been associated with diseases including cancer, amyotrophic lateral sclerosis (ALS), and frontal temporal dementia (FTD).^5–7^ A correlation has been found between an increased expression of hnRNP K and a lower chance of recovery in different cancers, including head and neck squamous cell carcinomas,^8^ prostate cancer,^9^ and hepatocellular carcinoma.^10^ Being a multifunctional protein, hnRNP K has been shown to be involved in regulating various oncogenes, including c-Myc^11^ and c-Src,^12^ that can lead to abnormal cell proliferation, metastasis, and resistance to cancer treatments.^13^

In ALS, lower expression of hnRNP K has been found to drive disease progression.^14,15^ hnRNP K is an important regulator of axogenesis, neuronal cell repair, and signal transduction, each of which is impaired in ALS patients.^16^ Although the actual onset of ALS is unclear, it has been shown that more than 30 hexanucleotide repeats (GGGGCC) are found in the *C9orf72* gene in 40% of the patients with familial ALS.^17,18^ The prevailing theory suggests that the hexanucleotide expansion gives rise to aberrant RNA transcripts that bind to RNA-binding proteins (RBPs), including hnRNP K, to form RNA foci, which are known to cause RNA toxicity.^17,19,20^ RNA foci can lead to inhibition of ribonucleotide reductase regulatory subunit M2 (RRM2) by binding to hnRNP K, and this ultimately interferes with the deoxyribonucleotides synthesis.^17^ Additionally, the translation of aberrant RNA transcripts can result in dipeptide repeats (DPR) such as polyglycine-alanine, poly-glycine-arginine, poly-proline-alanine, etc. These assemblies of aberrant transcripts and DPRs cause impairment in normal cellular regeneration, nerve signals, and DNA repair, which could eventually lead to ALS onset.^17,19,20^ A study conducted on the post-mortem motor cortex of *C9orf72* ALS patients demonstrated that hnRNP K deposits in the cytoplasm, causing the mis-localization of the protein and resulting in the dysregulation of RRM2 and DNA damage. Additionally, in a zebrafish model, increasing the levels of hnRNP K reduced the DNA damage caused by *C9orf72* RNA toxicity, thus suggesting a route to mitigate *C9orf72* ALS via increased hnRNP K expression.^7,21^

Given the abnormal expression of hnRNP K in multiple fatal and incurable diseases, it is important to understand the regulatory mechanisms of hnRNP K expression. While there are multiple ways to regulate gene expression either at the DNA or RNA level, translational control via the 3’-UTR offers flexibility by reversibly fine-tuning gene expression with minimal threats of permanent changes.^22^ The mRNA 3’-UTR is gaining attention due to its emerging roles in the modulation of post-transcriptional processing, including mRNA stability, translation, and degradation.^23,24^ Such translational control can be achieved via RNA binding proteins and microRNAs (miRNAs) interacting with the 3’-UTR.

miRNAs are short, non-coding RNAs about 22 nucleotides in length that repress translation by targeting the mRNA 3’-UTR using its complementarity miRNA seed sequence.^25^ The miRNA miR-1249-3p has been shown to target the *HNRNPK* 3’-UTR lowering the expression of hnRNP K under healthy physiological conditions.^26^ In diseased states, such as various cancers or ALS, the levels of both miR-1249-3p and hnRNP K are dysregulated, suggesting that this miRNA-mRNA regulation axis is more complicated than previously believed.^7,9,26^ Interestingly, we noted that the miR-1249-3p binding site within the *HNRNPK* 3’-UTR is guanine-rich, having the potential to fold into G-quadruplex (GQ) structures (Figure 1, top). G-rich DNA/RNA sequences with at least four guanine repeats can form Hoogsteen hydrogen bonds arranging themselves into guanine-quartet planes, which then stack with intercalated monovalent cations to form GQs.^27^ GQs have been shown to be important regulatory elements in transcription, mRNA processing, and stability.^28,29^ Moreover, they have been implicated in life-threatening diseases including cancer,^8,9,26,30^ neurological diseases^31^ such as fragile X syndrome,^32^ and ALS,^33^ and in viral systems like the human immunodeficiency virus-1 (HIV-1),^34^ SARS-CoV-2,^35^ Zika virus,^36^ and Ebola virus^37^. GQs have been shown to recruit RBPs, including the fragile X messenger ribonucleoprotein (FMRP) and fused in sarcoma (FUS) protein,^38,39^ as well as to interfere with miRNAs binding to their targets, highlighting roles of GQs in RNA metabolism.^40^ Studies have suggested that GQs forming in the 3’-UTR can regulate the accessibility of a miRNA binding site, either to expose it^41^ or to prevent the binding of an miRNA.^39^ While many GQs have been characterized in mRNA 3’-UTRs, studies about their involvement in regulating miRNA interactions with mRNA targets are still limited. In the case of the *HNRNPK* mRNA, the overlapping of a potential GQ-forming site with the miR-1249-3p binding site (Figure 1, bottom) suggests a possible competition between GQ structure formation and miR-1249-3p binding that could interfere with the miRNA-induced inhibition of the hnRNP K translation.

**Figure 1.**
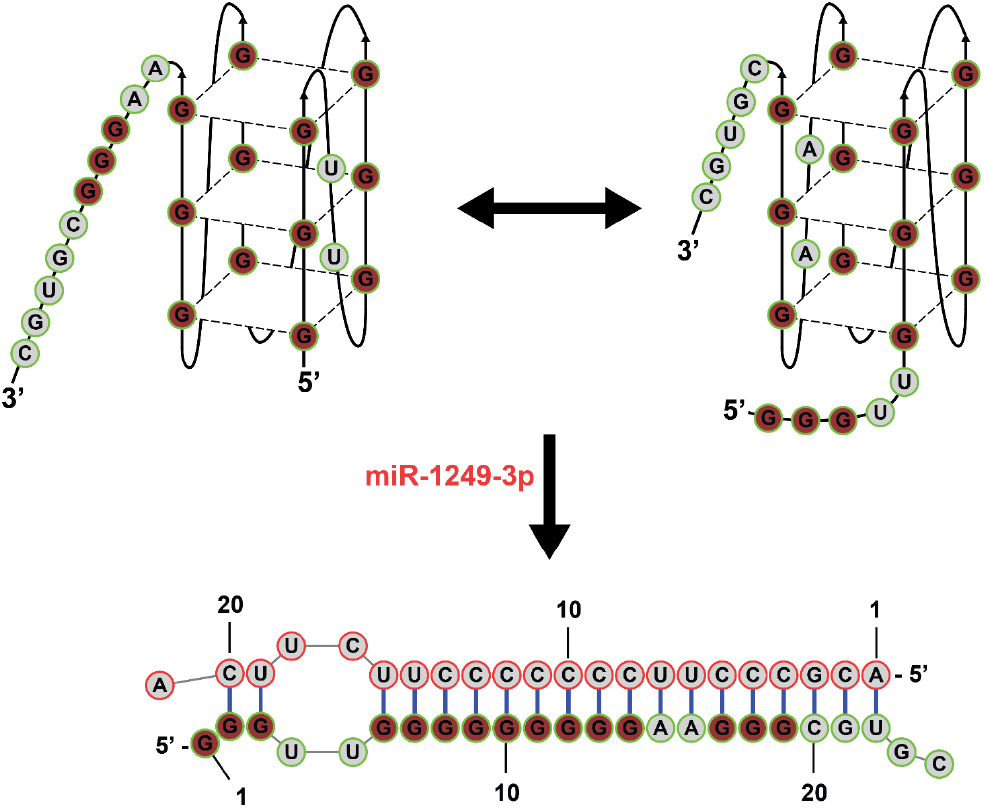
Five repeated G-tracts (maroon) involved in the GQ formation. G-tracts (1-4, top left) and G-tracts (2-5, top right) of *HNRNPK* arranged to form three-planar G-quartet structures. *HNRNPK* G-rich (GR) sequence (bold maroon, bottom) overlapped with the predicted miR-1249-3p binding site.

Thus, in this study we investigated the interactions between the *HNRNPK* 3’-UTR and miR-1249-3p to gain insight into the interplay between GQ and miRNA-mRNA duplex formation. Understanding the role of the GQ in modulating the accessibility of miR-1249-3p binding to the *HNRNPK* 3’-UTR will provide more clarity about the translational regulation of *HNRNPK,* potentially leading to the identification of novel therapeutic targets. We characterized the formation and potassium-dependent stability of a GQ in the *HNRNPK* 3’-UTR and demonstrated that the formation of this GQ structure modulates its binding interactions with miR-1249-3p *in vitro*. This was further supported by luciferase reporter assays, which demonstrated that the GQ could potentially act as a regulator of these binding interactions affecting *HNRNPK* translation.

## Results and Discussion

### The *HNRNPK* GR forms a stable intramolecular parallel GQ

We noted that the predicted binding site of miR-1249-3p within the *HNRNPK* 3’-UTR mRNA is G-rich, (sequence named here *HNRNPK* GR, Table 1), and using the QGRS mapper software^42^ we determined that it has the potential to form GQ structures. Thus, we employed various biophysical techniques to characterize the structure formed by *HNRNPK* GR. First, one dimensional (1D) proton ^1^H NMR spectroscopy was used to monitor the GQ signature imino proton resonance region in the 10-12 ppm range, which originates from the non-canonical base-pairing between adjacent G-G within the G tetrads.^43^ Imino proton resonances were present in the 10-12 ppm region, even in the absence of KCl, indicative of Hoogsteen base pairing in *HNRNPK* GR. Increasing concentrations of KCl resulted in some of the sharper resonances losing intensity with a concomitant increase in the intensity of upfield resonances, indicating that K^+^ ions stabilize a different GQ conformation than the one formed in their absence. The *HNRNPK* GR has 5 G-tracts available to form GQ structures, which require only four repeated G-tracts (Table 1). Thus, different GQ structures are possible (Figure 1) and the ^1^H NMR spectra suggest that the K^+^ ions are stabilizing a predominant GQ conformation. No resonances were observed in the region 12-14.5 ppm, corresponding to imino proton resonances of Gs and Us involved in Watson-Crick base, indicating that no hairpin formation is competing with the GQ structure formed by *HNRNPK* GR (Figure 2A).

**Table 1.**
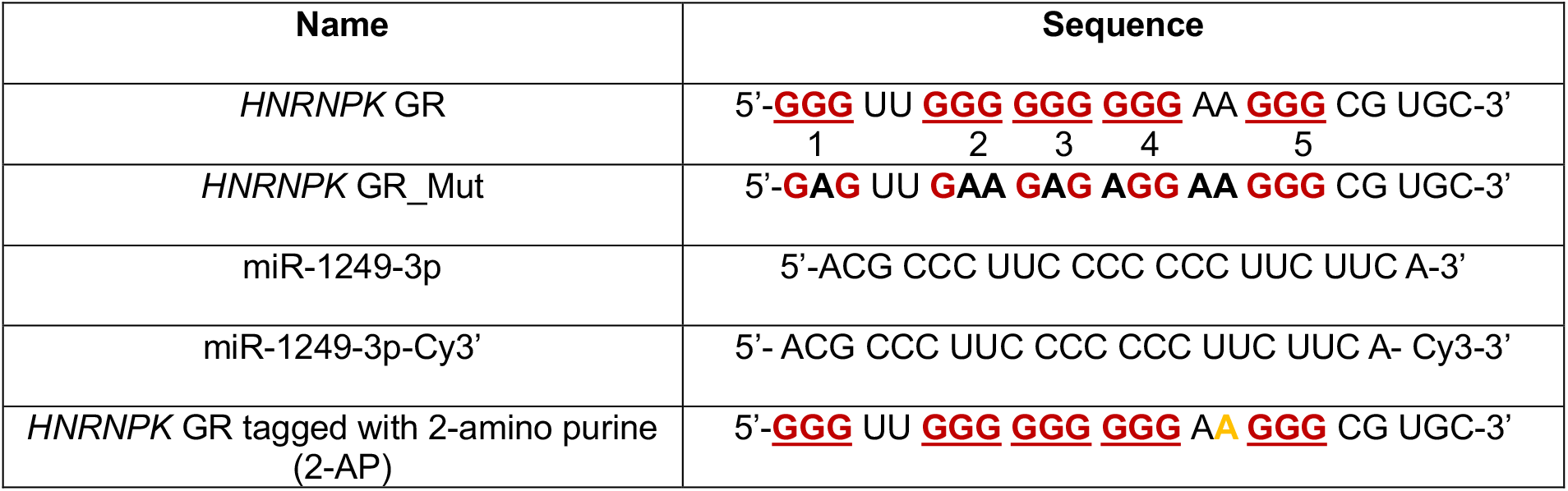
Sequences used in this study (Gs in the 3’-UTR of *HNRNPK* predicted to be involved in GQ formation are shown in red, mutated Gs in bold black, and 2-AP in bold yellow).

**Figure 2.**
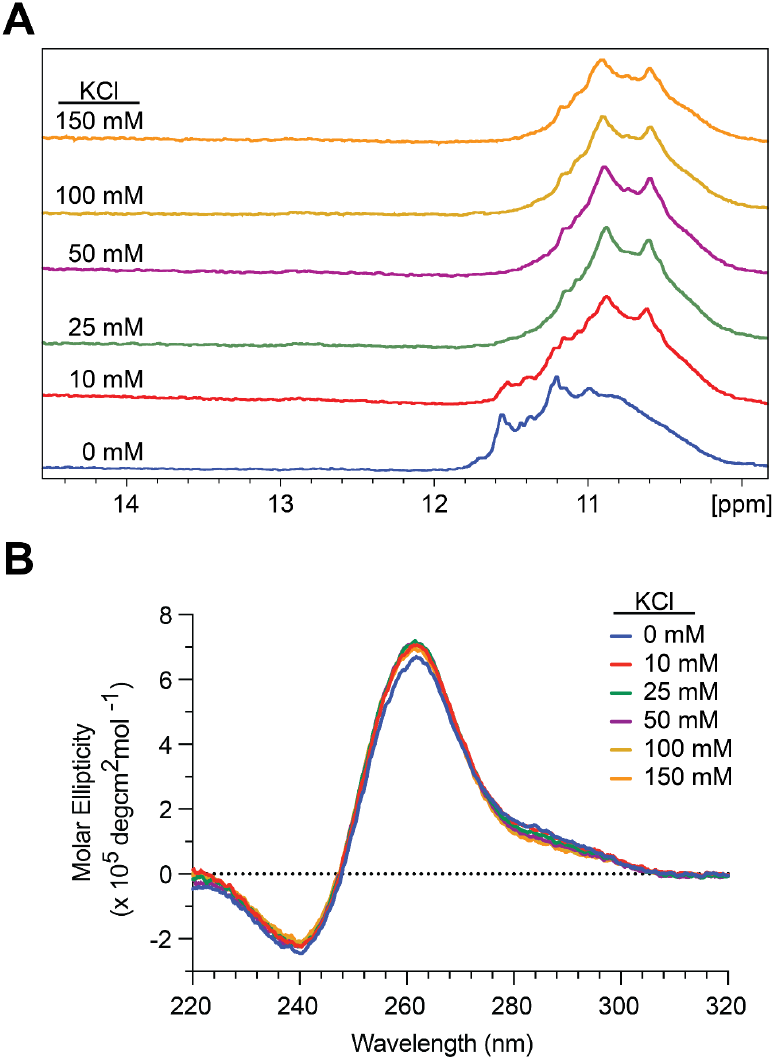
1D ^1^H NMR spectra (A) and CD spectra (B) of *HNRNPK* GR in the presence of increasing KCl concentrations.

Next, circular dichroism (CD) spectroscopy was employed to determine the topological conformation of the GQ formed by *HNRNPK* GR in the presence of increasing concentrations of KCl. In RNA, parallel GQs are favored due to the steric hindrance created by the ribose 2’-OH group, which allows the anti-conformation of the base rather than the syn-conformation.^44^ Parallel GQs give rise to a CD band pattern with a positive band at ∼265 nm and negative band at ∼240 nm, whereas anti-parallel GQs show a positive band at ∼295 nm and a negative band at ∼260 nm.^39,45^ The CD spectrum of *HNRNPK* GR in the absence of potassium has a positive maximum at ∼265 nm and negative minimum at ∼240 nm, confirming the formation of parallel GQs (Figure 2B). Minimal changes are observed upon the addition of K^+^, since the stabilization of a particular GQ structure by these ions observed in the 1D ^1^H NMR spectra cannot be captured by CD spectroscopy as any RNA GQ structure would be parallel giving rise to the same CD band signature.

Next, to evaluate the stability of the GQ, we performed ultraviolet light (UV) thermal denaturation spectroscopy experiments on 10 µM *HNRNPK* GR in the absence of K^+^ ions or in the presence of 150 mM KCl, monitoring the unfolding of the GQ structure at 295 nm, a wavelength sensitive to GQ dissociation with increasing temperature (Figure S1).^46^ In the absence of KCl, a single hypochromic transition was observed, which was fitted assuming a two-state model (eq 1, Materials and Methods) to determine a melting temperature (T_m_) of 52.3±0.1°C (Figure S1). The *HNRNPK* GR GQ was too stable to observe a full transition for its dissociation at 150 mM KCl (Figure S1). Thus, to determine the molecularity of the GQ structure, we carried out UV thermal denaturation experiments in 10 mM KCl varying the concentration of *HNRNPK* GR over a ten-fold range (Figure 3A). The T_m_ for 10 µM of *HNRNPK* GR at 10 mM KCl was determined to be 76.6±0.1°C (Figure 3A and eqs 1 and 2, Materials and Methods) and the enthalpy of GQ formation was determined to be -62.7±0.1 kcal/mol (eq 1, Materials and Methods). Since the enthalpy for a single G-tetrad was reported to be between -18 kcal/mol and -25 kcal/mol,^47^ these results suggest the formation of a three-plane GQ in *HNRNPK* GR.

**Figure 3.**
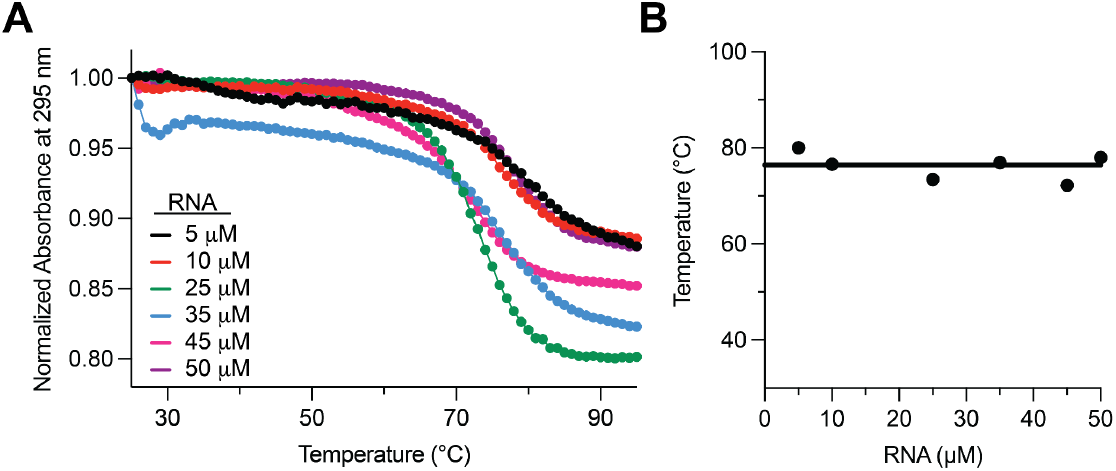
A) UV thermal denaturation spectroscopy of *HNRNPK* GR at different concentrations from 5-50 µM in 10 mM KCl. B) Plot of the *HNRNPK* GR melting temperature (T_m_) versus the RNA concentration.

At all RNA concentrations we observed hypochromic transitions (Figure 3A) where the T_m_ was not dependent upon the RNA concentration (Figure 3B, Table S1) indicating that *HNRNPK* GR forms an intramolecular GQ.^47^ In conclusion, we found that the GR region in the *HNRNPK* mRNA 3’-UTR forms an intramolecular parallel GQ structure stabilized by K^+^ ions.

### The *HNRNPK* GR GQ modulates miR-1249-3p binding

Previous studies found that miR-1249-3p binds to the 3’-UTR of *HNRNPK* to regulate protein expression,^26^ but the potential for a GQ structure to confer an additional level of regulation has not been explored. Therefore, we investigated the binding of miR-1249-3p to the *HNRNPK* GR, evaluating the influence of potassium ions, which stabilize GQ structures. 1D ^1^H NMR spectroscopy was carried out either in the absence or presence of 150 mM KCl, since at 0 mM KCl the *HNRNPK* GQ structure is not as stable as it is in 150 mM KCl (Figure S1).

No resonances were observed in the 1D ^1^H NMR of the free miR-1249-3p between 10-14.5 ppm, indicating the absence of any secondary structures (Figure 4A). The change from GQ imino proton resonances to Watson-Crick imino proton resonances allows the assessment of the binding interactions between miR-1249-3p and *HNRNPK* GR, since the GQ and the *HNRNPK* GR:miR-1249-3p duplex structures are mutually exclusive.

**Figure 4.**
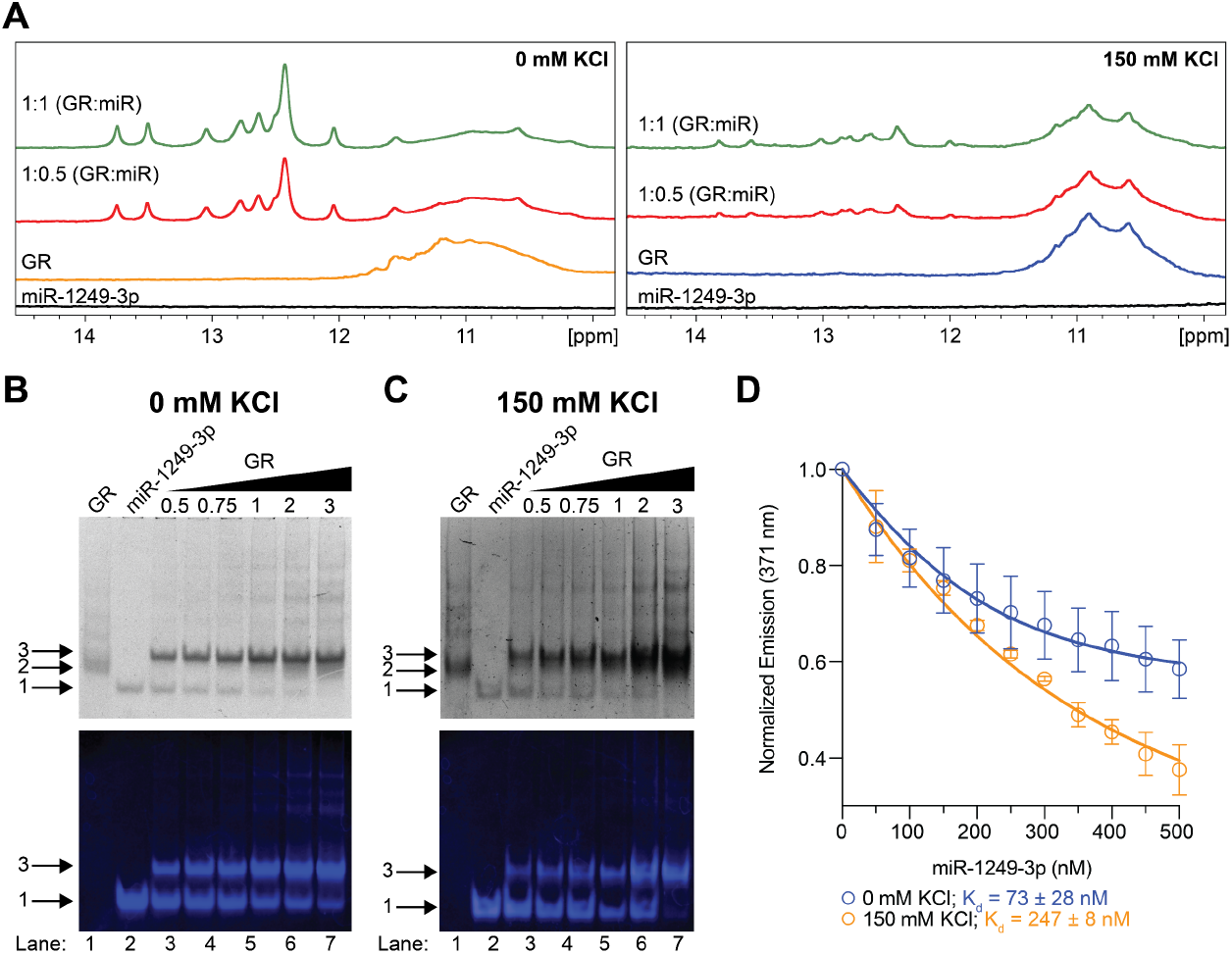
A) ^1^H NMR of the binding interaction between *HNRNPK* GR with miR-1249-3p at concentration ratios *HNRNPK* GR:miR (1:0.5 and 1:1) in the absence of KCl (left), and in 150 mM KCl (right). Native PAGE of the binding interactions between fluorescently labeled miR-1249-3p-Cy3 with *HNRNPK* GR at different concentration ratios ranging from 1:0.5 to 1:3 miR:*HNRNPK* GR in 0 mM KCl (B) and 150 mM KCl (C). D). K_d_ determination of the binding interaction between *HNRNPK* GR_2AP and miR-1249-3p: at 0 mM KCl and 150 mM KCl by fluorescence spectroscopy.

We initially titrated *HNRNPK* GR with miR-1249-3p in the absence of KCl, and observed the loss of resonance intensities between 10-12 ppm, suggesting a loss of Hoogsteen base pairing from the GQ structure,^43^ while simultaneously observing Watson-Crick imino proton resonances in the region 12-14.5 ppm^43^ corresponding to their complex formation (Figure 4A, left). When the experiment was repeated in 150 mM KCl, where the GQ structure is very stable, we still observed the appearance of new imino proton resonances between 12-14.5 ppm, but their intensity was much lower and there was only a slight decrease in intensity of the imino proton resonances between 10-12 ppm originating from the GQ structure (Figure 4A, right). Our previous results indicated the presence of the GQ in the *HNRNPK* GR in the absence of KCl, which is further stabilized by the K^+^ ions. Here, at 0 mM KCl the drastic loss of the Hoogsteen imino proton resonances indicates that although the *HNRNPK* GR adopts a GQ conformation, it is not stable enough to prevent the binding of miR-1249-3p.

However, in the presence of 150 mM KCl the loss of Hoogsteen imino proton resonances is less pronounced indicating that the stabilized GQ structure interfered with the binding of miR-1249-3p (Figure 4A, right). To further characterize the equilibrium between the *HNRNPK* GR GQ and *HNRNPK* GR:miR-1249-3p duplex, we incubated the *HNRNPK* GR:miR-1249-3p (1:1) for 7 days at 37°C and monitored complex formation (*HNRNPK* GR:miR-1249-3p) by acquiring 1D ^1^H NMR spectra at various times either in the absence or presence of 150 mM KCl. In the absence of KCl, we observed a further increase in the Watson-Crick base pairing with the concomitant disappearance of the GQ structure resonances (Figure S2, A). When the same experiments were repeated in 150 mM KCl (Figure S2, B), we observed a further increase in the Watson-Crick resonances, while the GQ structure resonances were still retained. These results indicate that the *HNRNPK* GR GQ and *HNRNPK* GR:miR-1249-3p duplex structures co-exist in equilibrium with one another and, once the GQ is stabilized by K^+^ ions, it affects the ability of miR-1249-3p to bind to *HNRNPK* GR.

To corroborate the 1D ^1^H NMR results, we used a fluorescently tagged miR-1249-3p-Cy3 to investigate the relative difference in its potassium dependent binding to *HNRNPK* GR by electromobility shift assay (EMSA). The gels were first imaged to observe the Cy3 fluorescence (Figure 4B and 4C, bottom), after which they were stained by SYBR Gold (Figure 4B and 4C, top). The free *HNRNPK* GR shows a main band (Figure 4B and 4C, top, lane 1, arrow 2) and additional upper bands corresponding to stacked GQ structures, which are not visible when observing the Cy-3 fluorescence (Figure 4B and 4C, top, lane 1). The Cy3 fluorescence of miR-1249-3p was used to observe the bound versus free miR in the native binding gel. The free miR-1239-3p-Cy3 (Figure 4B and 4C, lane 2, arrow 1), was shifted to a higher molecular weight complex when bound by *HNRNPK* GR (Figure 4B and 4C, lanes 3-7, arrow 3). As miR-1249-3p-Cy3 was incubated with increasing ratios of *HNRNPK* GR, in addition to the main band for the 1:1 miR-1249-3p-Cy3:*HNRNPK* GR complex, faint higher molecular weight fluorescent bands were also apparent (Figure 4B and 4C, bottom, lane 3-7). It is possible that these upper molecular weight complexes were formed by the higher order interactions between miR-1249-3p-Cy3:*HNRNPK* GR complexes. We observed that the intensity of the miR-1249-3p-Cy3:*HNRNPK* GR complex is higher at all *HNRNPK* GR concentrations ratios in 0 mM KCl as compared to 150 mM KCl (Figure 4B and 4C, bottom, lane 3-7). These results indicate that the binding interactions between *HNRNPK* GR and miR-1249-3p-Cy3 are weaker in 150 mM KCl as compared to 0 mM KCl, supporting our findings of a stable GQ interfering with miR-1249-3p binding (Figure 4B and 4C).

Finally, to obtain quantitative information about the binding interactions of miR-1249-3p with *HNRNPK* GR we employed steady state fluorescence spectroscopy, using a chemically modified *HNRNPK* GR with the adenine at position 16 in one of the GQ loops substituted for 2-amino purine (2-AP) (hereby named as *HNRNPK* GR_2-AP) (Figure 4D). We monitored the quenching of the 2-AP fluorescence due to the binding of miR-1249-3p at both 0 mM and 150 mM KCl to determine the dissociation constant (K_d_). The experiments were performed in triplicate, and the binding curves were fitted (eq 3, Materials and Methods) to determine the K_d_ for the binding interaction of miR-1249-3p with the *HNRNPK* GR_2-AP (73 ± 27 nM) in 0 mM KCl and (247 ± 8 nM) in150 mM KCl respectively. The increase in the K_d_ at 150 mM KCl compared to 0 mM KCl is consistent with the 1D ^1^H NMR spectroscopy and EMSA results showing the miR-1249-3p binding inhibition induced by the GQ stabilization due to the presence of K^+^ ions.

### The *HNRNPK* GR GQ regulates protein expression by suppressing translation

Given that our biophysical experiments indicated that the *HNRNPK* GR GQ stabilized by K^+^ ions reduced binding of miR-1249-3p *in vitro*, we analyzed its effect on protein expression in cells. Since the KCl concentration in cells is ∼ 140 mM, to destabilize/disrupt the GQ structure we mutated the *HNRNPK* GR sequence (*HNRNPK*

GR*_*Mut, Table 1). We carried out the biophysical characterization of *HNRNPK* GR*_*Mut by 1D ^1^H NMR and despite the mutations we observed imino proton resonances between 10-12 ppm, suggesting the formation of a GQ in this mutant (Figure S3, A). When mutating the G-tracts in the *HNRNPK* GR sequence, the G-tracts complementary to the seed sequence for miR-1249-3p were left intact (Table 1, tract 5) as they are essential for the miR binding to that region (Figure S3, D). The presence of this G-tract in *HNRNPK* GR*_*Mut together with the remaining Gs in tract 4 may contribute to the formation of this alternate GQ structure. To test the stability of this GQ, we analyzed *HNRNPK* GR_Mut by UV thermal denaturation spectroscopy and determined a melting temperature of 37.6±0.1°C in 150 mM KCl (Figure S3, B), indicating that it is significantly less stable than the wild-type *HNRNPK* GQ, which was too stable to fully dissociate at this KCl concentration (Figure S1). This result indicates that although *HNRNPK* GR_Mut forms a GQ *in vitro*, given its low melting temperature it is not stable enough to persist under physiological conditions.

Next, we carried out EMSA to investigate the binding interactions of the fluorescently tagged miR-1249-3p-Cy3 with the *HNRNPK* GR_Mut in 150 mM KCl. We observed the rapid depletion of free miR-1249-3p-Cy3 with the concomitant formation of the miR-1249-3p-Cy3:*HNRNPK* GR_Mut duplex (Figure S3, C, bottom, lane 3-7, arrows 1 and 3). These results suggest that although GQs are forming in *HNRNPK* GR_Mut, they are not stable enough to interfere with the miR binding interactions as all miR-1249-3p-Cy3 is complexed with *HNRNPK* GR when in a 1:1 ratio (Figure S3, C, lane 5).

Having successfully characterized GQ in the *HNRNPK* GR and binding interactions between *HNRNPK* GR with miR-1249-3p, we moved to luciferase assays in cells to investigate the regulatory role of *HNRNPK* GQ in the protein expression. We constructed reporter genes by cloning 367 nucleotides of the *HNRNPK* 3’-UTR, which includes the wildtype (GR) or mutant GQ sequence, downstream of the firefly luciferase gene in the pmiRGLO plasmid. These plasmids were then transfected into human hepatocyte derived carcinoma (Huh7) cells in which miR-1249-3p is endogenously expressed.^26^ After 48 h, we observed a significant reduction in the firefly luciferase activity for the mutant *HNRNPK* 3’-UTR (Figure 5A). This supports our *in vitro* data that the *HNRNPK* GQ interferes with the binding interactions of miR-1249-3p, as when the GQ is mutated miR-1249-3p can better access its binding site and repress expression.

**Figure 5.**
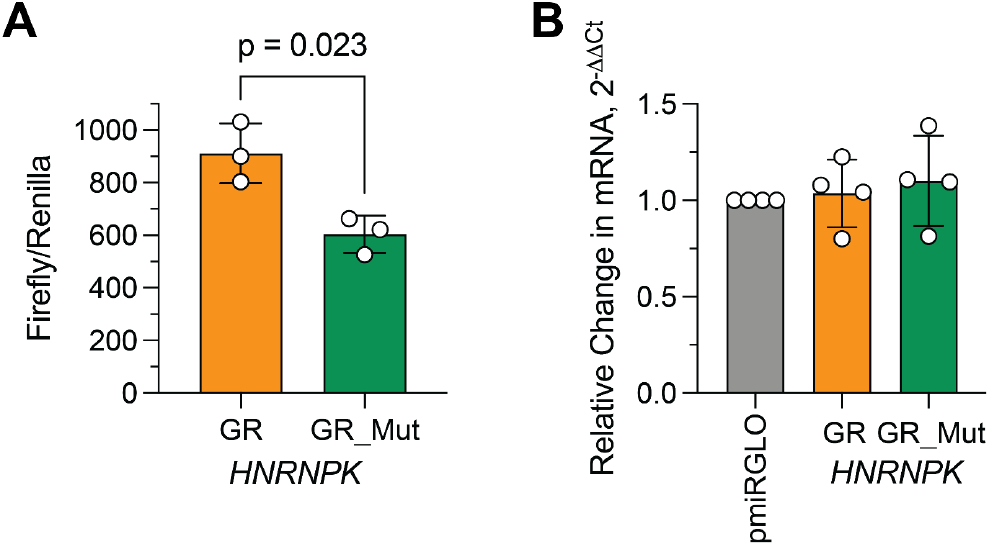
Role of the *HNRNPK* GR GQ on the firefly luciferase expression. A) A significant reduction in the relative luciferase activity in GR mut compared to GR. B) RT-qPCR showing no significant difference in the mRNA levels. Bars represent the mean and standard deviation of biological replicates (circles). The data represents mean and standard deviation of four biological replicates. Welch’s t-tests between WT and mutant were performed.

The observed reduction in luciferase expression could be affected by levels of mRNA transcripts produced by the DNA. To confirm that the difference in repression of the luciferase reporter observed between the wild type and mutant GR is not due to a difference in the transcription of the luciferase gene, we performed reverse transcription quantitative polymerase chain reaction (RT-qPCR) to determine the mRNA levels. We found that mRNA levels in Huh7 cells transfected with WT *HNRNPK* 3’-UTR or mutant *HNRNPK* 3’-UTR were not significantly different, indicating that the *HNRNPK* GQ does not have regulatory function at the DNA level (Figure 5B). The obtained result support a role of *HNRNPK* 3’-UTR GR in luciferase translation.

The mechanisms of post transcriptional regulation via secondary structures such as GQs are being explored but have not been leveraged to their full potential, especially regulation by RNA GQs.^44^ *In silico* analysis studies have predicted GQ formation in both the 5’-UTR and 3’-UTR of mRNA,^48,49^ but most of these studies have focused on GQs in the 5’-UTR, which have been shown to repress translation. For example, research conducted in human cell lines has demonstrated that expression of the *BCL-2* gene is downregulated due to the presence of a GQ in the 5’-UTR of its mRNA.^50^ Additionally, GQs characterized in the 5’-UTR of the *TRF2,*^51^ *Zic*-*1*,^52^ and *MT3*-*MMP*^53^ mRNAs have been shown to suppress the translation of these genes.

In comparison to the 5’-UTR, there has been limited exploration of the role of GQs in the 3’-UTR. The mRNA 3’-UTR plays an important role in RNA biogenesis and the presence of higher ordered structures, including GQs, has made this region an intriguing target for therapeutic study.^23,54^ For example, a study conducted by Crenshaw et al. demonstrated that amyloid precursor protein (APP) mRNA 3’-UTR contains GQs that play a role in decreasing protein expression.^55^ *PIM1*, a proto-oncogene found to be overexpressed in cancer, forms a GQ in the 3’-UTR of its mRNA which acts as a negative regulator of translation.^56^ In most cases, the presence of GQs in the 3’-UTR of mRNA results in lower translational efficiency. Additionally, the mRNA 3’-UTR is predominantly targeted by miRNAs which are an alternative mode of translational repression.^57^

There is a limited number of studies conducted investigating the role of GQs specifically in modulating the interactions of miRNAs which could alter translational regulation. In a study conducted by Rouleau et al., the miR-331-3p target site in the *FADS2* mRNA has G-rich sequence that folds into a GQ structure.^58^ This GQ in the target site was shown to interfere with miR-331-3p binding to the *FADS2* mRNA 3’-UTR, hence influencing its translation.^58^ Previously, *PSD-95* mRNA containing G-rich regions was shown to change between GQ conformations, allowing miR-125a to target *PSD-95* mRNA.^39^ In this study, we focused on the mRNA GQ-miRNAs axis for translational control of the hnRNP K protein. The results from our luciferase reporter assays suggest GQ-mediated regulation of miR-1249-3p binding to the 3’-UTR of the *HNRNPK* mRNA.

A single miRNA can target multiple genes and be involved in the regulation of multiple pathways. For example, miR-128-3p binds to 3′-UTR of the *VEGFC*,^59^ *VEGFR3*,^59^ and *PHF6*^60^ genes and results in decreased translation of each. Additionally, miR-128-3p has been shown to be involved in various pathways, including the Wnt/β-catenin,^61^ IRS-1/PI3K/AKT,^62^ and PI3K/Akt/NF-κB pathways.^63^ Similarly miR-1249-3p has been shown to involved in *HOXA13* ^64^ and *SKOR-1* regulation.^65^

Thus, targeting a miRNA to regulate the translation of a gene of interest could result in off-target effects of multiple biological pathways, an outcome that would be too complex to control. The presence of GQs in the target mRNA provides an additional layer of fine-tuned regulation of the gene of interest via either destabilizing or stabilizing its GQ structures. We found that mutating the *HNRNPK* GQ-forming sequence resulted in translational repression, suggesting that the GQ structure interferes with the binding interactions between miR-1249-3p and its targeted site in the 3’-UTR in cells. This suggests that the stability of the GQs in the *HNRNPK* mRNA governs the protein expression and controls the miR-1249-3p regulation of *HNRNPK*. However, we acknowledge the limitations of our study being conducted in hepatic cancer cells as compared to neuronal cells as hnRNP K is also involved in neurodegenerative disease .^5,7,21,26^

HnRNP K is implicated in the progression of multiple diseases, including cancer and ALS, but its contribution is context dependent – that is, the protein is overexpressed in some diseases but under-expressed in others.^7,66^ Our study provides crucial information about how the *HNRNPK* 3’-UTR GQ is involved in regulating protein expression and can be targeted for both high and low levels of hnRNP K. This allows for therapeutic design centered around the *HNRNPK* GQ to offset abnormal hnRNP K expression, making it a novel target for drug design.

## Materials and Methods

The *HNRNPK* mRNA sequence (NM_002140.5) was analyzed using the GQ predictive software QGRS Mapper,^42^ which identified a sequence within the *HNRNPK* 3’-UTR that has the potential to fold into a GQ (*HNRNPK* GR, Table 1). The *HNRNPK* GR, G-rich mutant sequence (*HNRNPK* GR_Mut), 2-amino purine fluorescently labeled *HNRNPK* GR variant (*HNRNPK* GR_2AP), miR-1249-3p and its fluorophore labeled variant miR-1249-3p-Cy3 which contains the Cy3 fluorophore at its 3’ end, were purchased from Horizon Discovery (Table 1).

### One-Dimensional Proton Nuclear Magnetic Resonance Spectroscopy

All 1D ^1^H NMR spectroscopy experiments were conducted using a 500 MHz Bruker NMR spectrometer (Topspin 3.2 software), with the water suppression achieved by using the WATERGATE water pulse sequence. All samples were prepared using a 225 µM RNA sample in 10 mM cacodylic acid pH 6.5 with 10% D_2_O at 20^°^C at the indicated concentrations of KCl.

To characterize the binding of miR-1249-3p, 1D ^1^H NMR experiments were carried out in two conditions: (i) in the absence of KCl and (ii) in the presence of 150 mM KCl. In both cases, *HNRNPK* GR samples were analyzed with snap-cooled miR-1249-3p at the indicated ratios of *HNRNPK* GR:miR-1249-3p. Time-dependent spectra were acquired after incubation at 37°C for the indicated times. Free miR-1249-3p was prepared at 225 µM, as described above, in control experiments.

### Circular Dichroism Spectroscopy

All CD experiments were performed on a Jasco J-810 spectropolarimeter at 25°C, scanning samples from 220 nm to 320 nm with a bandwidth of 2 nm, a 1 second response time, and a 1 mm path length. All data were acquired averaging seven scans. The GQ conformation in *HNRNPK* GR was investigated using a 20 µM sample in 10 mM cacodylic acid pH 6.5 in the presence of the indicated KCl concentrations. All data was corrected with a 10 mM cacodylic acid pH 6.5 blank.

### UV Thermal Denaturation Spectroscopy

A Cary Series UV-vis Spectrophotometer (Agilent Technologies) was used to investigate the stability of the structures formed by *HNRNPK* GR and *HNRNPK* GR_Mut. 10 µM *HNRNPK* GR was prepared in cacodylic acid (10 mM, pH 6.5), boiled for 5 minutes, and cooled at room temperature at the indicated concentrations of KCl. The change in absorbance was monitored at 295 nm from 25°C to 95°C at a rate of 0.2°C/min. The same procedure was repeated for preparing 30 µM *HNRNPK* GR_Mut sample in 150 mM KCl. Similarly, samples at the indicated concentrations of *HNRNPK* GR in 10 mM KCl were prepared in cacodylic acid (10 mM, pH 6.5), boiled for 5 minutes, and cooled at room temperature.

The thermodynamic parameters for *HNRNPK* GR and *HNRNPK* GR_Mut were determined by fitting the UV thermal denaturation curve to eq 1, which assumes a two-state model: ^67,68^

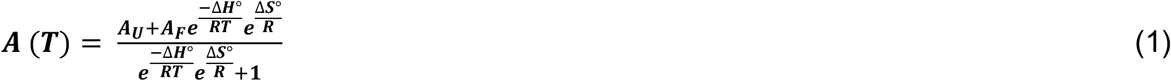

where A_U_ represents the absorbance of unfolded RNA, A_F_ represents the absorbance of folded RNA, R is the universal gas constant. The melting temperature (T_m_) was determined using eq 2:

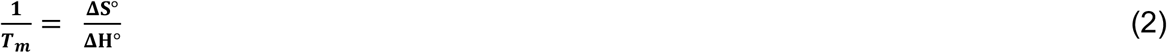

### Electromobility Shift Assay

To characterize the binding interactions of Cy3-labeled miR-1249-3p (miR-1249-3p-Cy3) with *HNRNPK* GR, 1 µM of snap-cooled miR-1249-3p was incubated for 30 minutes with indicated concentration of *HNRNPK* GR and KCl at room temperature. The samples were run on a 15% native polyacrylamide gel (30:0.8 acrylamide:bisacrylamide) for 4 hours (75 V, 4°C) in ½x tris boric acid EDTA (TBE) buffer. The gels were visualized in a LI-COR *Odyssey* Fc at 600 nm to observe the Cy3 fluorescence, followed by staining with SYBR® Gold to visualize all the RNA bands. The experiments were carried out in triplicate.

### Steady State Fluorescence Spectroscopy

All fluorescence spectroscopy experiments were conducted using a Horiba Jobin Yvon Fluoromax-4 fluorimeter with a 150 W ozone-free xenon arc lamp. Samples at 200 nM were prepared in 10 mM cacodylic acid, pH 6.5, in 0 mM or 150 mM KCl. The excitation wavelength was set to 310 nm with the bandpass excitation and emission of monochromators adjusted to 5 nm and the emission spectra were recorded at 371 nm (25°C). miR-1249-3p was titrated in 50 nM increments to the *HNRNPK* GR_2-AP sample in a 3 mm path-length quartz cuvette (Starna cells) in the presence of a 5-fold excess of a basic peptide derived from the capsid protein of the hepatitis C virus.^69^ All values were normalized to the free *HNRNPK* GR_2-AP fluorescence intensity at 371 nm. Determination of the dissociation constant (K_d_) was performed by fitting the normalized fluorescence data of each experiment to eq 3:^67^

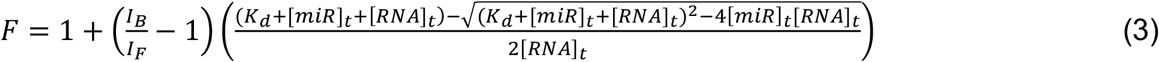

I_F_ and I_B_ denote the steady-state fluorescence intensities of free and bound RNA, respectively. [RNA]_t_ is the constant total RNA concentration, while [miR]_t_ is the total miR-1249-3p concentration. The reported K_d_ value is the average of three independent experiments with the standard error.

### Molecular Cloning

gBlocks of the *HNRNPK* 3’-UTR (NM_002140.5, nts:1616-1982) containing the *HNRNPK* GR and *HNRNPK* GR_Mut sequences were purchased from Integrated DNA Technologies. *HNRNPK* GR and *HNRNPK* GR_Mut were cloned into the pmiRGLO plasmid (Promega) by using HiFi Assembly (New England Biolabs) between the Nhe1Kos and Xba1 restriction sites. Generated sequences were named as pmiRGLO-*HNRNPK* GR and pmiRGLO-*HNRNPK* GR_Mut respectively. All plasmid sequences were confirmed using whole plasmid sequencing (Eurofins).

### Dual Luciferase Assay

Huh7 cells (gift from John Schoggins, University of Texas Southwestern Medical School) were cultured in DMEM (Gibco) supplemented with 10% fetal bovine serum (FBS; Corning Incorporated) and 100 units/mL penicillin and 100 μg/mL streptomycin sulfate (MP Biomedicals) at 37°C and 5% carbon dioxide.

On day 0, Huh7 cells were seeded in a 96-well white plate at 3×10^4^ cells/well. On day 1, cells were transfected with 300 ng of pmiRGLO-*HNRNPK* GR or pmiRGLO-*HNRNPK* GR_Mut in 50 µL of DMEM medium using DharmaFECT (Horizon Discovery) as per manufacturer’s instruction. After 48 h, the Nano-Glo Dual Luciferase Reporter Assay system (Promega) was used to measure the firefly and *Renilla* luciferase activities on a SpectraMax ID3 instrument (Molecular Devices). Samples were normalized by dividing the firefly by *Renilla* luminescence.

### Reverse Transcription Quantitative Polymerase Chain Reaction

Huh7 cells were plated in 6-well plate (Costar) at a density of 4×10^5^ cells/well on day 0. The following day, cells were transfected with 1 µg of pmiRGLO-3’-UTR, pmiRGLO-*HNRNPK* GR, or *pmiRGLO*-*HNRNPK* GR_Mut in 2 ml of DMEM using DharmaFECT as per manufacturer’s instruction. After 48 h, total RNA was extracted as described previously^70^ using TRIzol (Thermo Fischer) as per manufacturer’s instruction, and then treated with DNase I (Invitrogen). cDNA was prepared by reverse transcribing 1 µg of RNA (SuperScript IV Reverse Transcriptase; Thermo Fisher) using 1 µl of 50 mM random hexamer primers (Invitrogen). The obtained cDNA was then diluted 1:5 with nuclease-free water (Qiagen). 1 µl of diluted cDNA was added to 5 µl of SYBR Green (Invitrogen), 1 µl of 10 µM primer pairs (Table 2), and 3 µl of nuclease-free water. 40 cycles of amplification were performed using a QuantStudio 3 Real-Time PCR System (Applied Biosystems). Transcript levels were determined using the ΔΔCt and normalized to pmiRGLO-3’-UTR for each RNA.

**Table 2.**
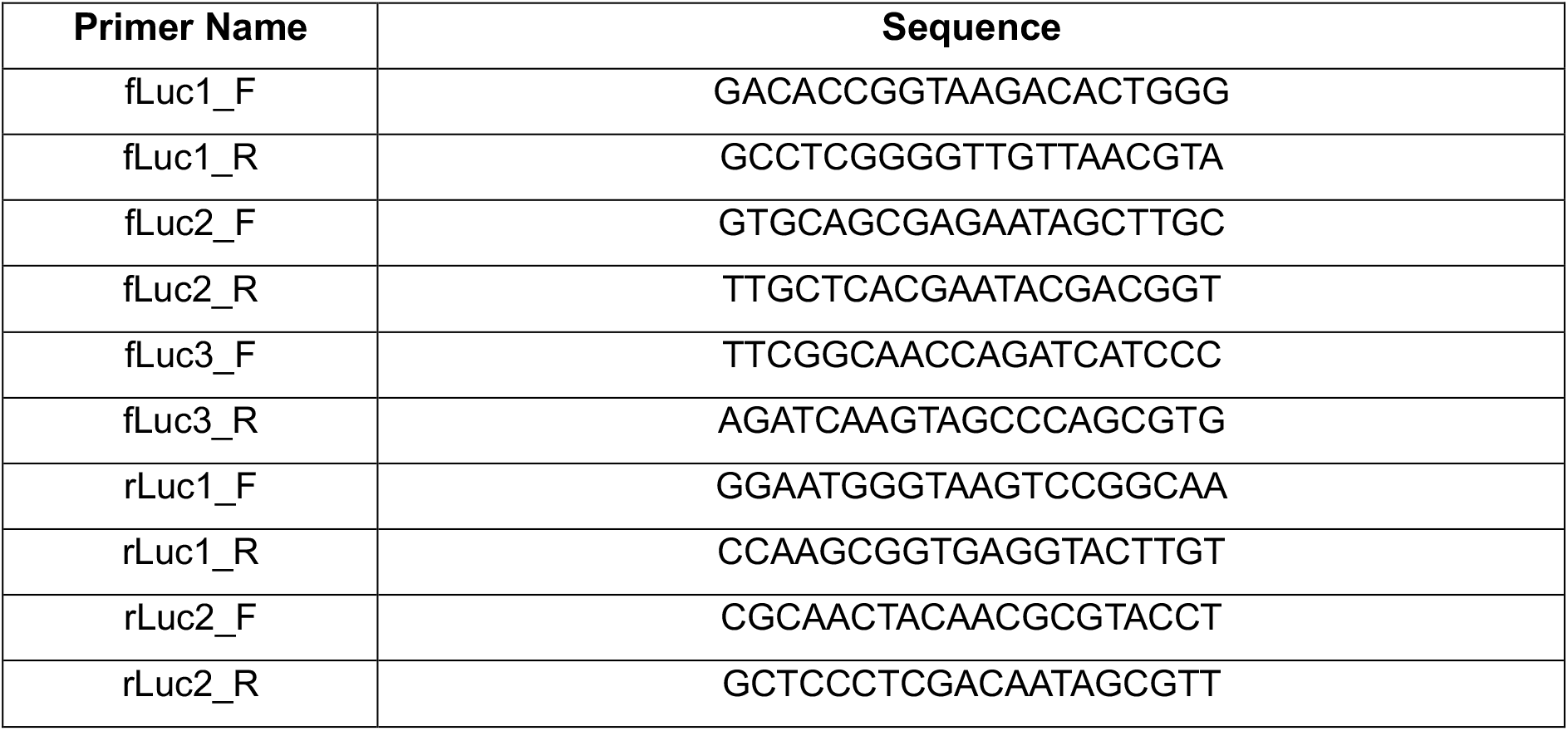
Primer sequences used for *HNRNPK* Reverse Transcription Quantitative Polymerase Chain Reaction.

## Author Contributions

MRM and DBH conceived the presented idea and supervised the work. MK, RMP, CG, LG, and JS performed the experiments and data analysis. MK and RMP wrote the initial manuscript and worked with DBH and MRM for preparation of final manuscript.

## Acknowledgements

This work was supported by the National Institute of General Medical Sciences 2R15GM127307-05, and the National Institute of Neurological Disorders and Stroke 5R25NS100118-07.

## Supporting Information

UV thermal denaturation spectroscopy of *HNRNPK* GR at 0 mM and 150 mM KCl (Figure S1); 1D ^1^H NMR of the binding interaction between *HNRNPK* GR and miR-1249-3p at a 1:1 concentration ratio of *HNRNPK* GR:miR; 1D ^1^H *HNRNPK* GR:miR complex incubated at 37°C for 2 days, 4 days, and 7 days in 0 mM and 150 mM KCl (Figure S2); 1D ^1^H NMR spectroscopy of the *HNRNPK* GR_Mut at 0 mM and 150 mM KCl (Figure S3, A); UV thermal denaturation of 30 µM *HNRNPK* GR_Mut at 150 mM KCl (Figure S3, B); native PAGE of the binding interactions between the fluorescently labeled miR-1249-3p-Cy3 with *HNRNPK* GR_Mut at different concentration ratios ranging from 1:0.5 to 1:3 in 150 mM KCl (Figure S3, C); The predicted base-pairing between *HNRNPK* GR and miR-1249-3p using RNAstructure software (Figure S3, D); melting temperatures of *HNRNPK* GR as a function of the RNA concentration (Table S1).

