## Supplemental figures for "The role of a G-quadruplex structure in mediating the interactions of miR-1249-3p with the 3’-UTR of the *HNRNPK* mRNA"

**Supporting Information**

**Content**

***Supplemental***

***Figures…………………………………………………………………………………………..1***

**Figure S1………………………………………………………………………………………..1**

**Figure S2………………………………………………………………………………………..1**

**Figure S3………………………………………………………………………………………..2**

***Supplemental Tables***

**Table S1. Thermodynamic parameters for *HNRNPK* GR GQ………………………….3**

**Supplementary Figures**

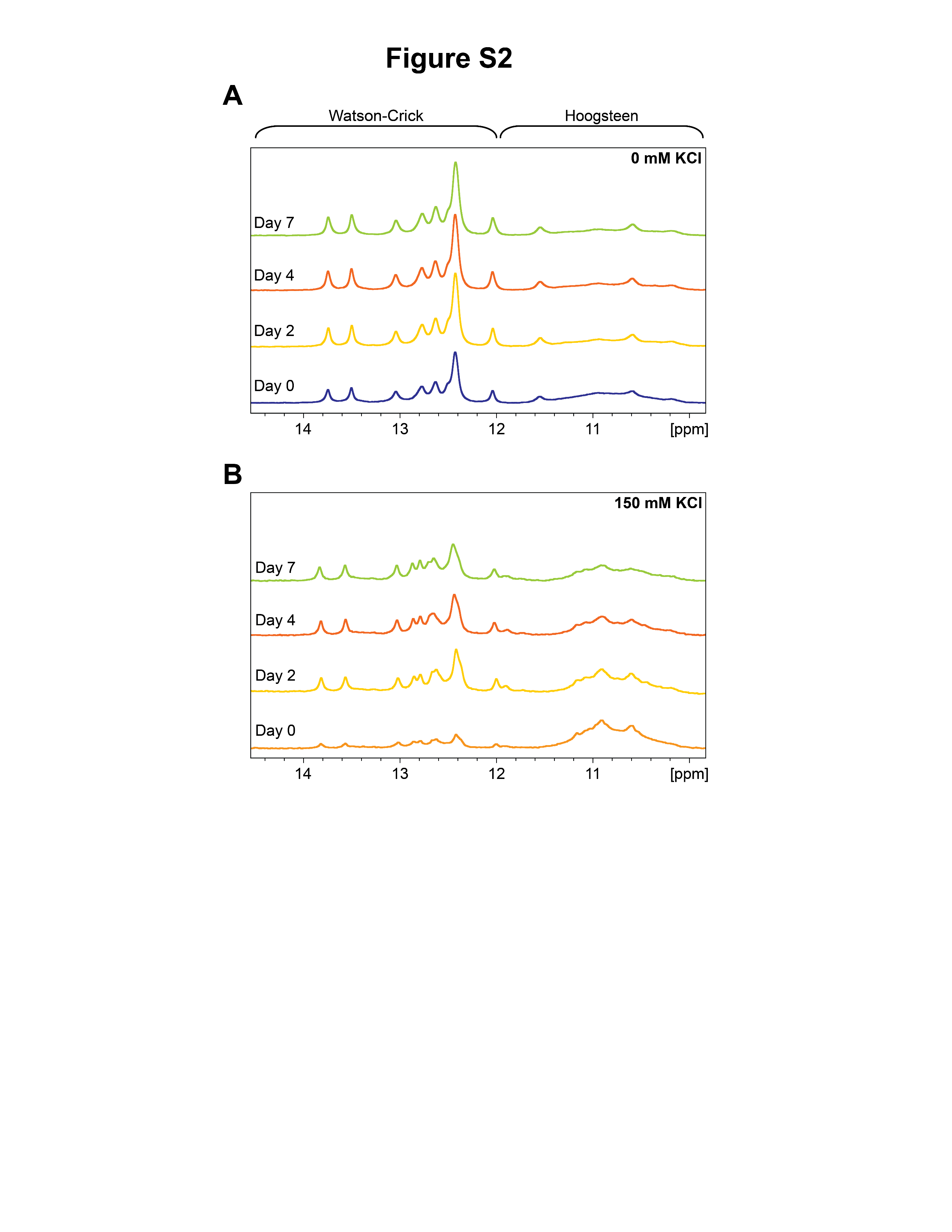

**Figure S2**. 1D ^1^H NMR of the binding interaction between *HNRNPK* GR with miR-1249-3p at a 1:1 concentration ratio of *HNRNPK* GR:miR. *HNRNPK* GR:miR complex incubated at 37°C for 2 days, 4 days, and 7 days in 0 mM KCl (top, A) and 150 mM KCl (bottom, B).

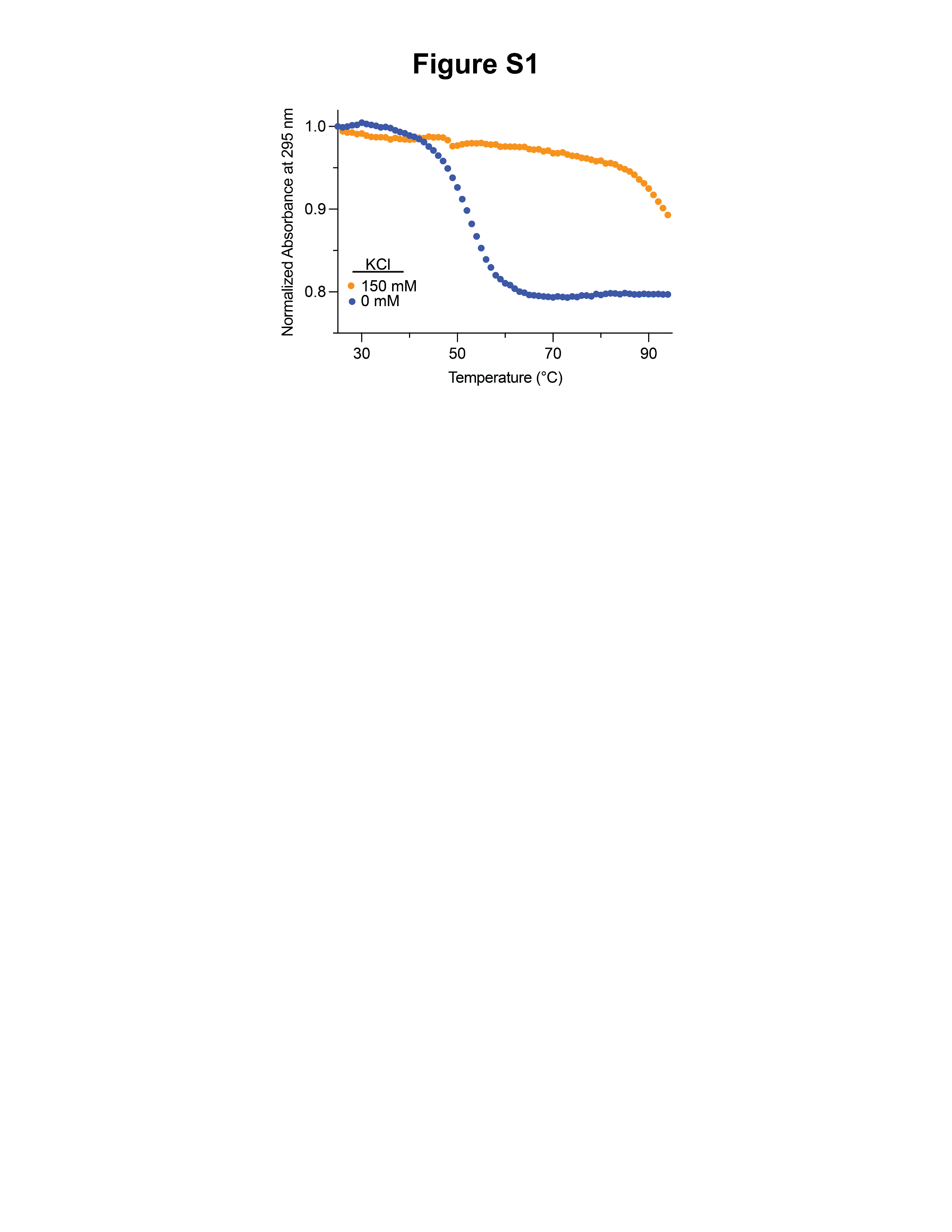

**Figure S1.** UV thermal denaturation spectroscopy of *HNRNPK* GR at 0 mM (blue) indicating GQ dissociation via hypochromic transition. At 150 mM KCl (orange), the GQ was too stable to observe a full hypochromic transition.

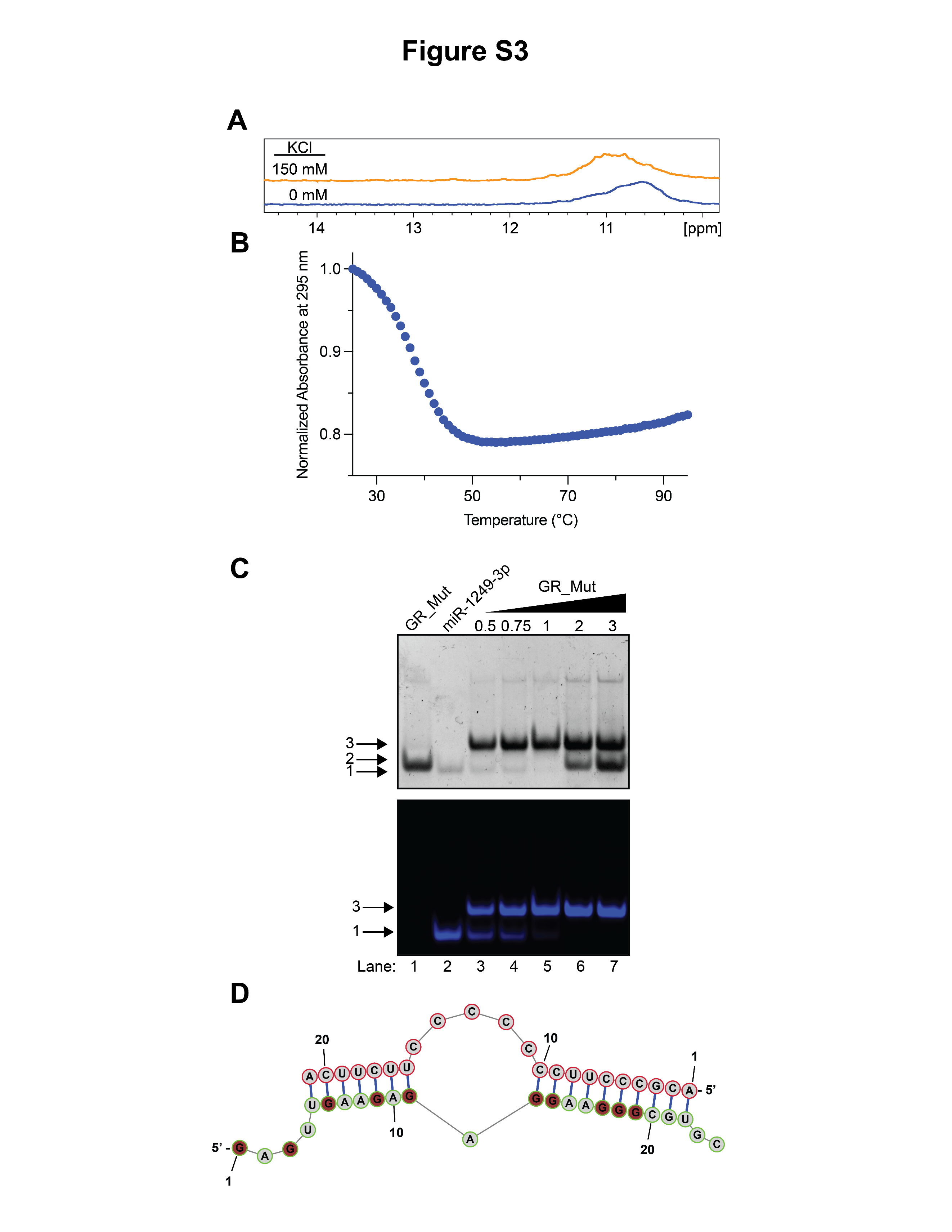

**Figure S3**. A) 1D ^1^H NMR spectroscopy of the *HNRNPK* GR_Mut at 0 mM and 150 mM KCl. B) UV thermal denaturation of 30 µM *HNRNPK* GR_Mut at 150 mM KCl. C) Native PAGE of the binding interactions between the fluorescently labeled miR-1249-3p-Cy3 with *HNRNPK* GR_Mut at different concentration ratios ranging from 1:0.5 to 1:3 miR:*HNRNPK* GR_Mut in 150 mM KCl. D) The predicted base-pairing between *HNRNPK* GR and miR-1249-3p using RNAstructure.

**Supplemental Tables**

**Table S1. Melting temperature of *HNRNPK* GR GQ as a function of the RNA concentration**

| **RNA (µM)** | **T_m_ (°C)** |
| --- | --- |
| 5 | 80.0±0.1 |
| 10 | 76.6 ±0.1 |
| 25 | 73.4±0.1 |
| 35 | 76.9±0.1 |
| 45 | 72.2±0.1 |
| 50 | 78.0±0.1 |
